# Ultra-accurate Microbial Amplicon Sequencing Directly from Complex Samples with Synthetic Long Reads

**DOI:** 10.1101/2020.07.07.192286

**Authors:** Benjamin J Callahan, Dmitry Grinevich, Siddhartha Thakur, Michael A Balamotis, Tuval Ben Yehezkel

## Abstract

Out of the many pathogenic bacterial species that are known, only a fraction are readily identifiable directly from a complex microbial community using standard next generation DNA sequencing technology. Long-read sequencing offers the potential to identify a wider range of species and to differentiate between strains within a species, but attaining sufficient accuracy in complex metagenomes remains a challenge. Here, we describe and analytically validate LoopSeq, a commercially-available synthetic long-read (SLR) sequencing technology that generates highly-accurate long reads from standard short reads. LoopSeq reads are sufficiently long and accurate to identify microbial genes and species directly from complex samples. LoopSeq applied to full-length 16S rRNA genes from known strains in a microbial community perfectly recovered the full diversity of full-length exact sequence variants in a known microbial community. Full-length LoopSeq reads had a per-base error rate of 0.005%, which exceeds the accuracy reported for other long-read sequencing technologies. 18S-ITS and genomic sequencing of fungal and bacterial isolates confirmed that LoopSeq sequencing maintains that accuracy for reads up to 6 kilobases in length. Analysis of rinsate from retail meat samples demonstrated that LoopSeq full-length 16S rRNA synthetic long-reads could accurately classify organisms down to the species level, and could differentiate between different strains within species identified by the CDC as potential foodborne pathogens. The order-of-magnitude improvement in both length and accuracy over standard Illumina amplicon sequencing achieved with LoopSeq enables accurate species-level and strain identification from complex and low-biomass microbiome samples. The ability to generate accurate and long microbiome sequencing reads using standard short read sequencers will accelerate the building of quality microbial sequence databases and removes a significant hurdle on the path to precision microbial genomics.

## Introduction

In recent years, the development of long-read sequencing technologies and concomitant advances in their cost-efficiency and accuracy have brought disruptive change to a variety of important biological applications. For example: Long-read sequencing has largely trivialized the generation of complete and accurate de novo bacterial genomes (Koren 2015), has expanded and improved the enumeration of transcriptional isoforms (Wang 2016; Beiki 2019; Wang 2020) and immune repertoires (Brochu 2020), and has vastly improved the detection and description of structural genetic variation (Mahmoud 2019).

The characterization of bacterial species and strains directly from complex microbial samples using amplicon sequencing — in which PCR-amplified DNA fragments (amplicons) from complex genetic mixtures are sequenced — is still an ongoing challenge in microbiology in part due to short sequencing reads not containing enough information to support highly resolved phylogenetic classification calls. Long-read sequencing has become an increasingly attractive technology for amplicon sequencing (Callahan 2019; Kumar 2019; Johnson 2019). However, while long-read amplicon sequencing approaches based on PacBio and Oxford Nanopore sequencing technologies are being developed and deployed in a wide variety of applications (Caskey 2017; Earl 2018; Lam 2020), a combination of error rates, cost and more limited availability of long-read sequencing capacity continue to impede their widespread application.

Synthetic long-read (SLR) sequencing technologies are appealing because they can leverage inexpensive, accurate, and widely available short-read sequencing platforms such as those from Illumina to generate accurate long-read sequencing data that would be otherwise harder or impossible to obtain. However, SLR technologies that were previously commercialized by 10x Genomics (Zheng 2016) and Moleculo (Kuleshov 2014) were not compatible with amplicon sequencing because they assign the same DNA identifier to multiple molecules in the same well/droplet, which is not amenable to reconstructing the sequence of single long molecules. Other SLR methods exist that utilize unique molecular identifiers (UMIs) instead of well/droplet identifiers to tag each DNA molecule with a DNA identifier that can be later read by DNA sequencing to identify each molecule, but their chemistries limit their read lengths (Burke 2016, Karst 2018) and these methods have not been commercialized.

This is the first evaluation of a commercially-available long read microbiome sequencing technology that builds upon previous developments in SLR sequencing (Stapleton 2016; Hong 2014) to address the challenge of accurately identifying species and strains directly from complex microbial samples using only a standard short read DNA sequencer. The synthetic long-read sequencing developed by Loop Genomics employs unique molecular identifiers that enable the reconstruction of single, long molecules from short sequencing reads. If this technology truly enables single-molecule long read sequencing of mixtures of highly homologous molecules, it would be an attractive option for a multitude of long-read amplicon sequencing applications. Loop Genomics has recently commercialized synthetic long-read amplicon sequencing under the LoopSeq product line which is available both as a kit and a laboratory service (Loop Genomics, CA). However, there have been no peer-reviewed evaluations of the recently commercialized technology to date.

In this paper, we report on the exceptional accuracy of LoopSeq SLR technology on a defined mixture of known bacterial sequences (a mock community) and develop guidelines for filtering and processing LoopSeq SLR sequencing data. We compare LoopSeq full-length 16S rRNA gene amplicon sequencing results to the current gold standard of PacBio CCS sequencing on a common set of human fecal microbiome samples. After denoising, the overall community compositions measured by LoopSeq and PacBio CCS from the same fecal samples were highly concordant. However, LoopSeq achieved higher levels of long-read amplicon sequencing accuracy, and that higher accuracy was likely maintained in complex communities based on the frequencies with which inferred differences between gene variants occurred in the conserved or variable regions of the 16S rRNA gene. Finally, we show how LoopSeq full-length 16S sequencing can be used to identify CDC defined foodborne pathogen species from samples of US retail meat and to distinguish distinct strains within those species.

## Results

### Accuracy and error modes: 16S sequencing of the Zymo mock community

The ZymoBIOMICS Microbial Community DNA Standard (the Zymo mock community) consists of genomic DNA from 8 bacterial strains of the species *Bacillus subtilis, Enterococcus faecalis, Escherichia col i, Lactobacillus fermentum, Listeria monocytogenes, Pseudomonas aeruginosa, Salmonella enterica*, and *Staphylococcus aureus.* We used the LoopSeq 16S Long Read Kit (Loop Genomics, CA) to barcode and amplify the full-length 16S rRNA gene (Methods), which was then sequenced by Loop Genomics using Illumina NextSeq500 PE150. The assembled long reads were filtered to remove those that did not contain both primers, that had lengths outside the expected range (1400-1600 nts) or that contained more than two expected errors according to their quality scores (Edgar 2015). The ~83% of reads that passed filtering were processed by the DADA2 method using default parameters (Methods) to produce a set of denoised amplicon sequence variants (ASVs) discriminated at single-nucleotide resolution ( Callahan 2017).

All of the 27 denoised ASVs from the Zymo 16S rRNA data were determined to represent true 16S rRNA gene alleles present in the mock community strains (Figure S1) using the same evaluation approach previously described for PacBio long-read amplicon sequencing data (Callahan 2019). The full complement of intragenomic variation present in the multiple alleles of the 16S rRNA gene within each mock community strain was recovered along with the correct ratio of the unique gene copies within each strain’s genome. No contaminant sequences (i.e. sequences originating from outside the mock community) and no false positive sequences (i.e. sequences containing uncorrected errors) were present in the denoised ASVs.

LoopSeq long amplicon sequencing reads were highly accurate. 94.6% of these ~1500nt reads contained no errors at all, a substantially higher fraction than the ~50-80% error-free short-reads (100-300nts) produced by standard Illumina amplicon sequencing. The DADA2 denoising method (Callahan 2016) was used to associate error-containing LoopSeq reads to the true sequence from which they most likely originated, and the locations, type and quality score associated with each point error were recorded (Methods, Figure 1). Insertion and deletion errors were extremely rare (<2 x 10^-6^ per nucleotide) and there was no evidence of specific read positions associated with significantly higher insertion or deletion error rates. Substitution errors were somewhat more common (4.6 x 10^-5^ per nucleotide) and occurred at a slightly higher rate near the start and end of the reads. This was predicted by lower quality scores and expected from the lower coverage of the long-read contigs by the short-reads at the ends of the contigs. Overall, this per-nucleotide error rate of ~5 x 10^-5^ per nucleotide, alternatively expressed as a 99.995% per-nucleotide accuracy, significantly exceeds the best accuracy results currently reported in the literature for amplicon sequencing using standard Illumina sequencing or Pacific Biosciences CCS long-read sequencing (Pfeiffer 2018; Callahan 2019).

**Figure 1:**
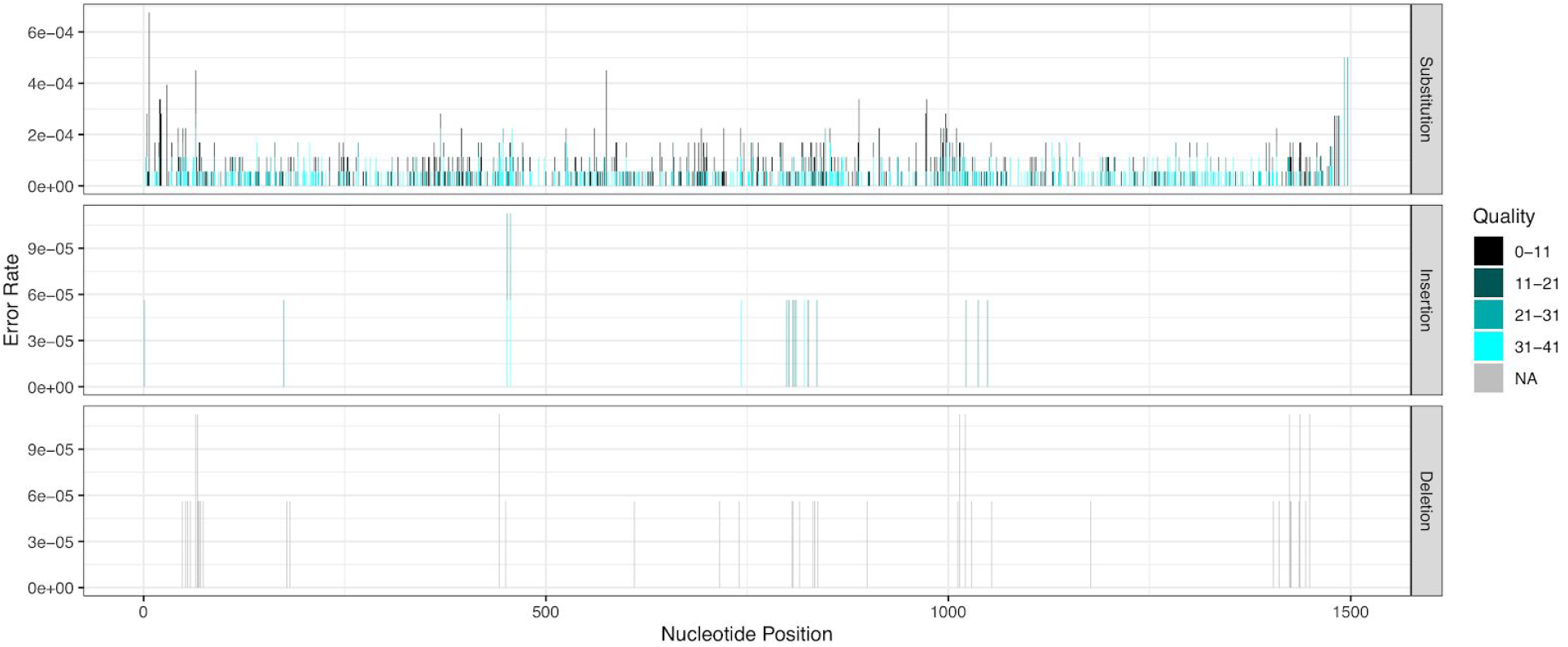
The rates of LoopSeq point errors by position in read, type (in/del/substitution) and quality score.

We performed a further manual inspection of potential structural errors in LoopSeq amplicon sequencing data, for example PCR chimeras that are formed through processes that modify large segments of the sequencing read rather than a single position at a time. We identified a type of structural error we refer to as an introgression, in which a segment of one amplicon is replaced (or is introgressed into) the homologous segment in another amplicon (Figure 2a). This error mode arises due to an interaction between PCR chimeras and the assembly of short-read into an SLR. The presence of early-round PCR chimeras can result in a segment of chimeric DNA being selected by the assembler when constructing the long-read sequence from all of the short-reads sharing that UMI. Usefully, lower quality scores are typically found in the introgressed segment, reflecting the lower level of short-read consensus at those positions.

**Figure 2:**
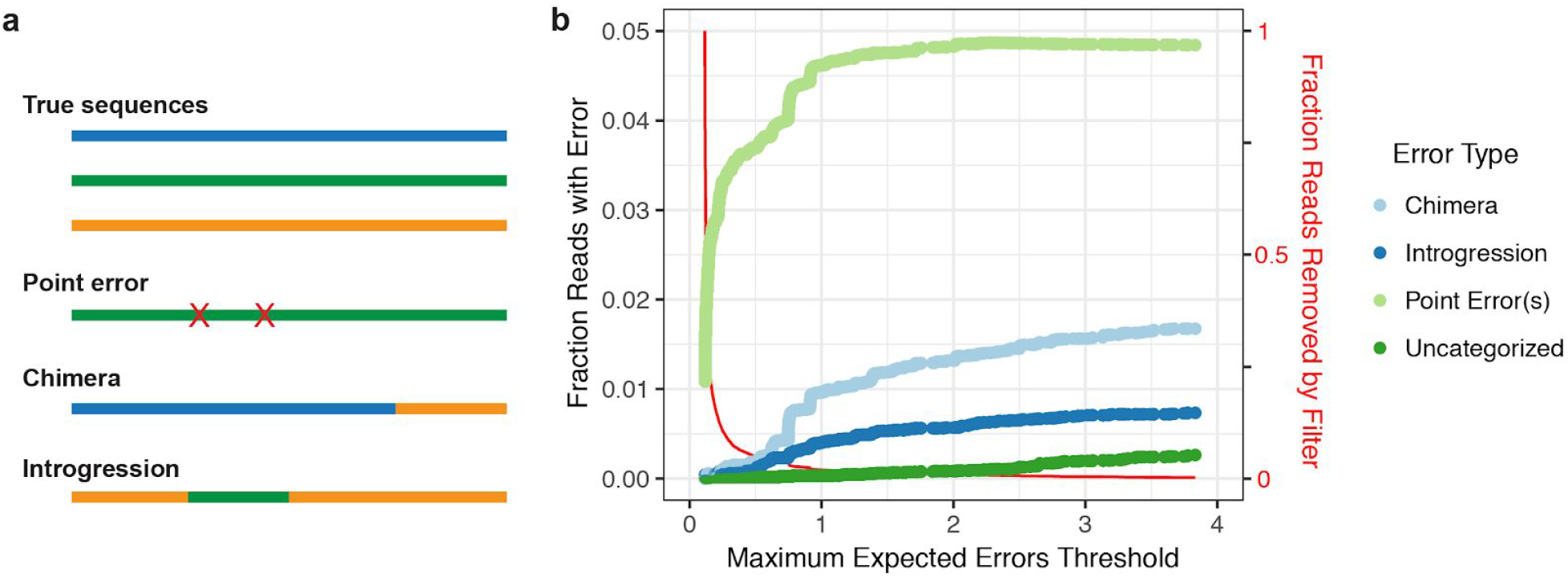
Error types and rates in LoopSeq data. a) Schematic description of three error types: point errors, chimeras, introgressions. b) The fraction of each error type as a function of expected error quality filter threshold. The red line shows the fraction of LoopSeq reads removed as a function of the filter threshold.

We developed a moving window approach to identify structural errors, both typical PCR chimeras and introgressions, in the uncorrected LoopSeq reads from this mock community of known composition (Methods). For each read, we determined whether it was correct (i.e. contained no errors) or incorrect. If it was incorrect, we determined whether it was a chimera, an introgression, or contained point errors. The fraction of reads of each of these error types is plotted as a function of the expected errors filtering threshold (Edgar 2017) for the unfiltered LoopSeq reads from the Zymo mock community in Figure 2b. A small number of incorrect long reads could not be unambiguously categorized, perhaps because they were structural errors that also contained point errors.

Structural errors (chimeras and introgressions) accounted for just over 2% of the reads in the unfiltered LoopSeq amplicon reads from the Zymo mock community, but this fraction significantly decreased with stricter quality filtering (Figure 2b). For this sequencing library, a threshold of 0.5 maximum expected errors appeared to effectively balance the removal of reads by the filter (~3%) with the suppression of structural errors (<0.5%). We explored the effects that this filtering has on high-sensitivity sample inference with the DADA2 method (Methods). We found that more stringent filtering allowed singleton detection (DETECT_SINGLETONS=TRUE) and a much more sensitive ASV detection threshold (OMEGA_A=1e-10) to be accurately applied to LoopSeq data. Only 4 false positive ASVs were identified each represented by a single read, and that collectively accounted for <0.03% of all denoised reads. New methods for *de novo* identification of introgressions could largely eliminate those remaining rare false positive ASVs and allow for long-read amplicon sequencing with single-nucleotide resolution, a singleton frequency threshold, and a near-zero false positive rate.

### Accuracy of longer reads: 18S-ITS and genomic sequencing of fungal and bacterial isolates

We performed LoopSeq amplicon sequencing of the approximately 2.3 kilobase 18S-ITS gene region from isolates of six fungal species obtained from the ATCC: *Saccharomyces cerevisiae, Aspergillus oryzae*, *Candida albicans, Trichoderma reesei, Kluyveromyces lactis, Penicillium chrysogenum*. LoopSeq reads were filtered for the presence of the forward and reverse primers and then trimmed to the region between the primers. In each sample, over 80% of the reads had identical sequences that also exactly matched the 18S-ITS gene region from a previously sequenced isolate of that fungal species. We determined these reads to be the error-free fraction of the data (Figure 3). This may be a slight underestimate if some reads with different sequences represented error-free reads from minority alleles of the many (100s) of copies of the 18S-ITS gene region present in some fungi.

**Figure 3:**
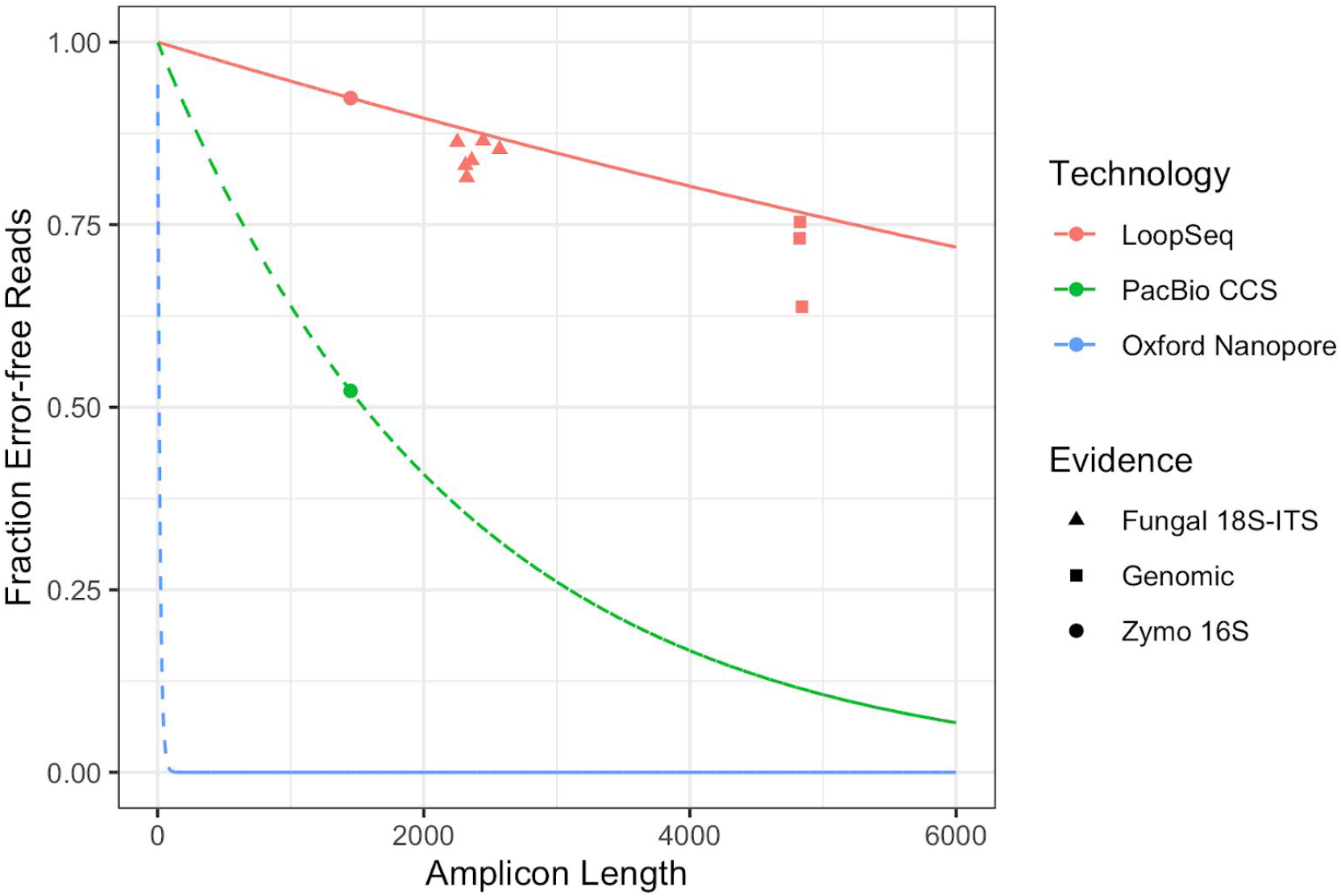
The fraction of error-free amplicon sequencing reads of different lengths using commercially available long-read sequencing technologies. Points represent observations from measurements of defined communities (the Zymo mock community) or single-strain isolates (fungal and bacterial isolates) that were reported in this manuscript for LoopSeq, and that were reported in Callahan 2019 for PacBio CCS reads. The 16S rRNA and fungal 18S-ITS data is from traditional amplicon sequencing of a single genetic region (Methods). The LoopSeq genomic amplicon data is based on LoopSeq sequencing of randomly amplified regions of the genome of several bacteria (Methods). Lines represent the expected error-free fraction based on measurements of the error rates in 16S rRNA gene amplicon sequencing data, and assuming a constant per-base error rate. The Oxford Nanopore line is based on the per-base error-rate of 6% reported by the manufacturer for the R10 chemistry.

We also performed LoopSeq sequencing of randomly amplified segments of the genomes from isolates of three bacterial species obtained from the ATCC: *Nitrosomonas europaea, Desulfovibrio desulfuricans*, and *Salinispora tropica.* LoopSeq reads were filtered for length between 4000 and 6000 bases, resulting in a median read length of ~5Kb. The error-free fraction was determined to be those reads that exactly matched the associated reference genome. The error-free fraction of these LoopSeq reads was over 60% in all three species (Figure 3). This may be a slight underestimate if errors exist in the reference genomes, or if non-genomic elements such as plasmids were present in the sequencing data.

### Performance in complex communities: Human fecal samples

We performed LoopSeq full-length 16S amplicon sequencing of the DNA extracted from three human fecal samples. The same extracted DNA had been previously characterized by PacBio full-length 16S amplicon sequencing (Callahan 2019). The raw LoopSeq data was filtered and processed by the DADA2 method using default parameters (Methods). The ASV tables produced by LoopSeq and by PacBio (two replicates, using two versions of their Sequel chemistries) were merged into a common ASV table.

The communities measured by LoopSeq and PacBio were highly concordant. A median of 89.9% of the reads detected by LoopSeq in each sample shared the same sequence with reads also detected by PacBio, while the same measure between the PacBio replicates was 94.1%.

We used the Bray-Curtis metric to quantify community-wide dissimilarity between the measured communities in each sample by each technology. Visualization of those results in a PCoA ordination plot (Figure 4) revealed that differences between LoopSeq and PacBio measurements of the same sample were trivial compared to differences between samples. In fact, the median Bray-Curtis dissimilarity between LoopSeq and PacBio measurements of the same sample was just 0.187, barely higher than the median 0.173 Bray-Curtis dissimilarity between replicate PacBio measurements of the same sample.

**Figure 4:**
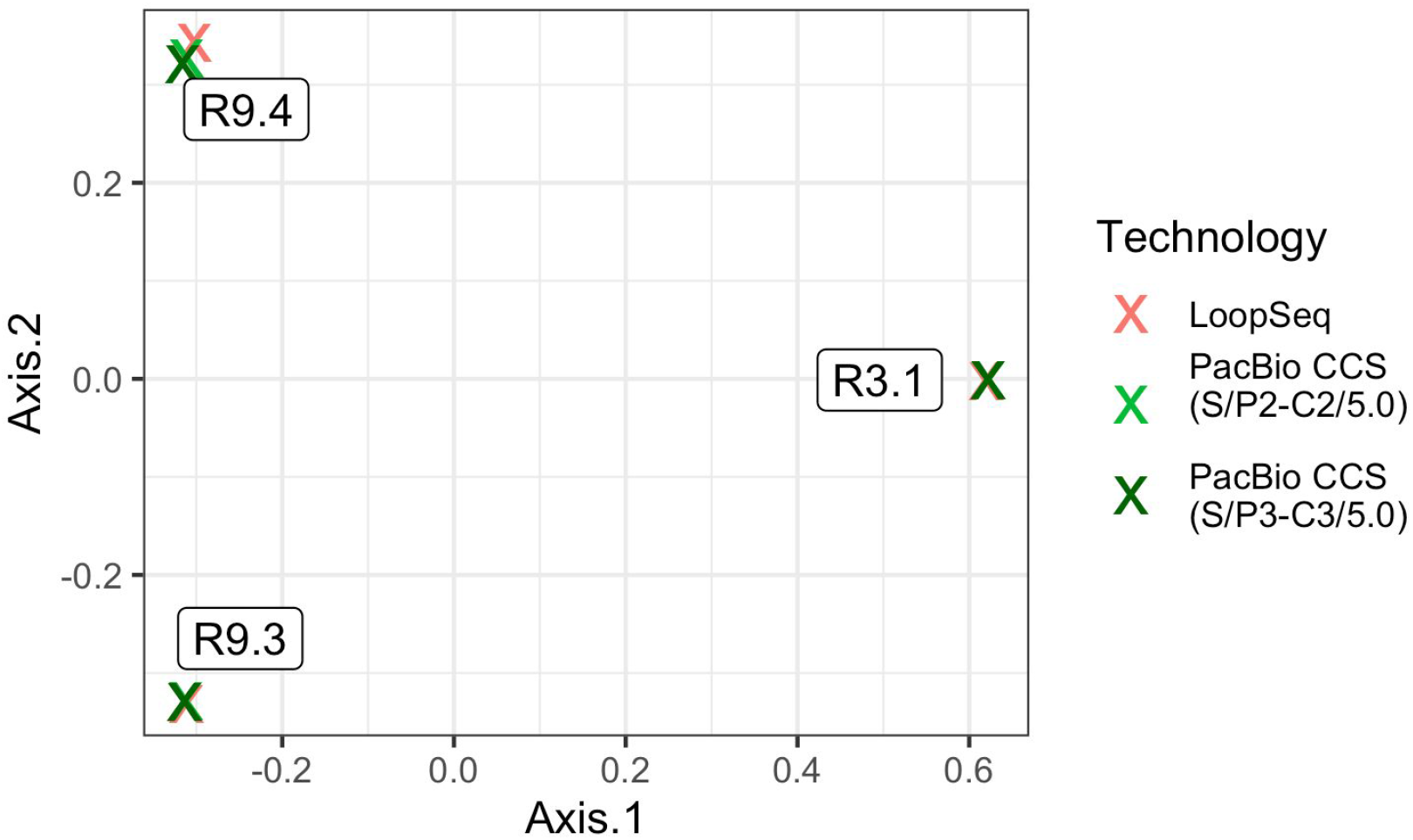
PCoA ordination of the total community compositions of three human fecal samples as measured by LoopSeq and PacBio full-length 16S rRNA gene sequencing. The community compositions measured by each technology were highly similar, leading to the data points on this ordination being highly overlapping for each sample. PacBio CCS measurements were made using two different sequencing chemistries as indicated in the legend parentheticals.

The differences between LoopSeq ASVs were highly enriched at known variable positions of the 16S rRNA gene, supporting high LoopSeq accuracy in the human fecal samples. We performed high-sensitivity sample inference on these human fecal samples using the DADA2 method (Methods) which also provides a full description of the substitutions between each ASV and the “sibling” ASV from which it was distinguished by the denoising algorithm. We used the ssu-align program (Methods) to define whether substitutions between sibling ASVs occurred at conserved or variable regions of the 16S rRNA gene (Nawrocki 2009). The results of this analysis for each ASV identified by DADA2 in high-sensitivity mode are shown in Figure 5.

**Figure 5:**
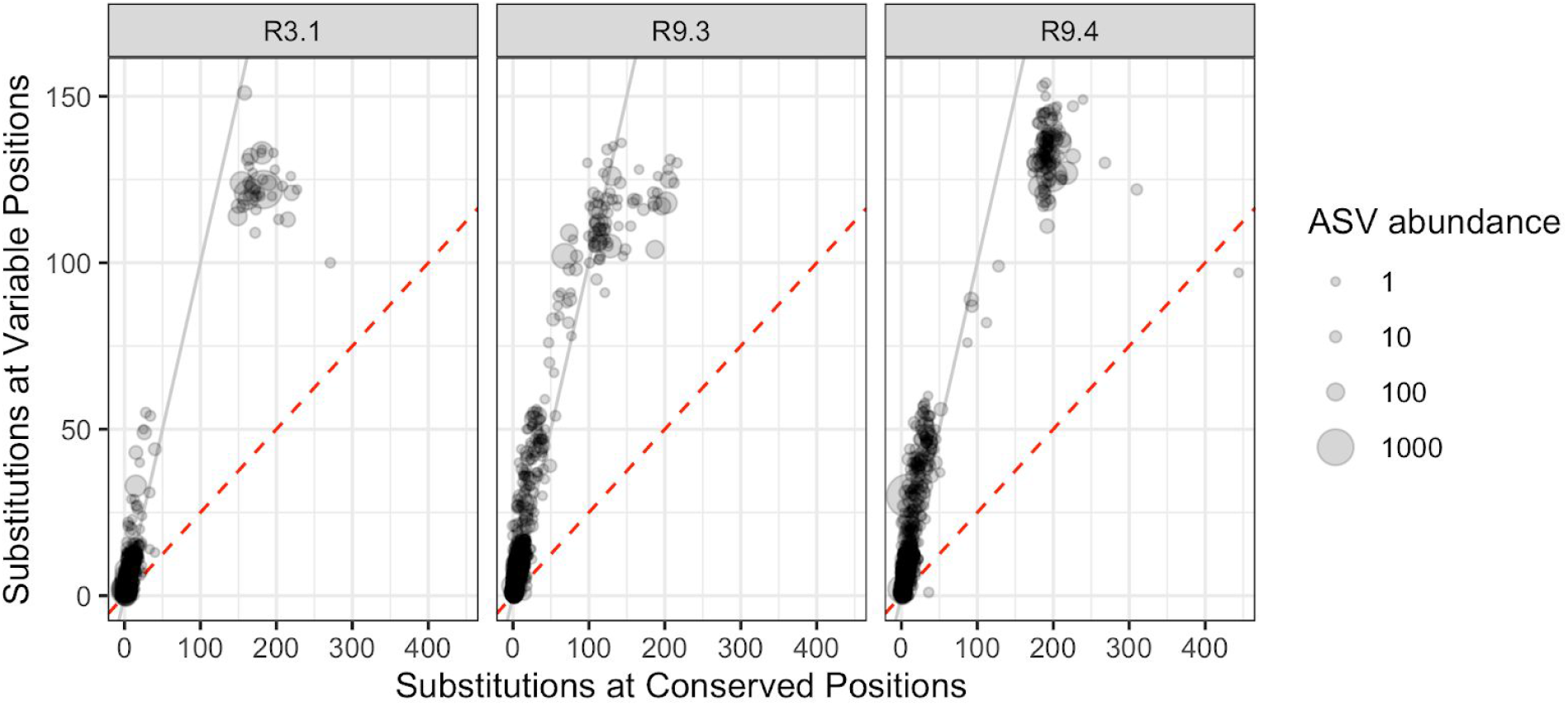
The number of substitution differences between DADA2-denoised ASVs and the next closest ASV that are in conserved vs. variable regions of the 16S gene, in LoopSeq data from three human fecal samples. Approximately 80% of nucleotide positions are conserved, thus if substitution patterns were random (as might be the case if they were caused by sequencing errors) then points should fall on the dashed red line.

Substitutions between sibling LoopSeq ASVs occur with approximately equal frequency in variable positions of the 16S rRNA gene and in conserved regions. This is far in excess of what would be the case if substitutions were distributed randomly given that ~80% of the 16S rRNA gene nucleotides are conserved (Methods). This represents about a 4-fold enrichment in substitutions at variable sites relative to the null expectation of randomly located substitutions from sequencing errors. Overall, this pattern supports the high accuracy of LoopSeq long-read amplicon sequencing in complex community samples, consistent with the results on the simpler mock community.

### Example application: Foodborne pathogens in retail meat

To investigate the potential for ultra-accurate long-read 16S sequencing to identify and track foodborne pathogens, we performed LoopSeq 16S sequencing on DNA extracted from a rinsate of six samples of US retail meat (Methods). We were particularly interested in the foodborne pathogen species that the CDC has identified as of particular importance in retail meat: *Yersinia enterocolitica*, *Escherichia coli, Salmonella enterica, Clostridium perfringens, Campylobacter spp.*, and *Listeria monocytogenes.* We performed high-sensitivity sample inference on these retail meat samples using DADA2, and assigned taxonomy down to the genus level using the naive Bayesian classifier method and the Silva database (Methods). Denoised ASVs with genus assignments that matched high-interest foodborne pathogens were then given species assignments if their BLAST results unambiguously supported a particular species (Methods).

The accuracy and length of the full-length LoopSeq 16S sequences allowed us to unambiguously identify all of the foodborne pathogen species of interest, four of which were present in these samples (Figure 6a). *Y. enterocolitica* was clearly distinguishable from other closely related *Yersinia* in these samples that are not typically pathogens in humans, including the notorious metagenomic false positive *Y. pestis* (Afshinnekoo 2015a; Ackelsberg 2015; Afshinnekoo 2015b). *C. perfringens* was distinguishable from the closely related *C. septicum,* and we were able to accurately identify the *Salmonella* strain in sample GT5 as *S. enterica* subsp. *enterica.*

**Figure 6:**
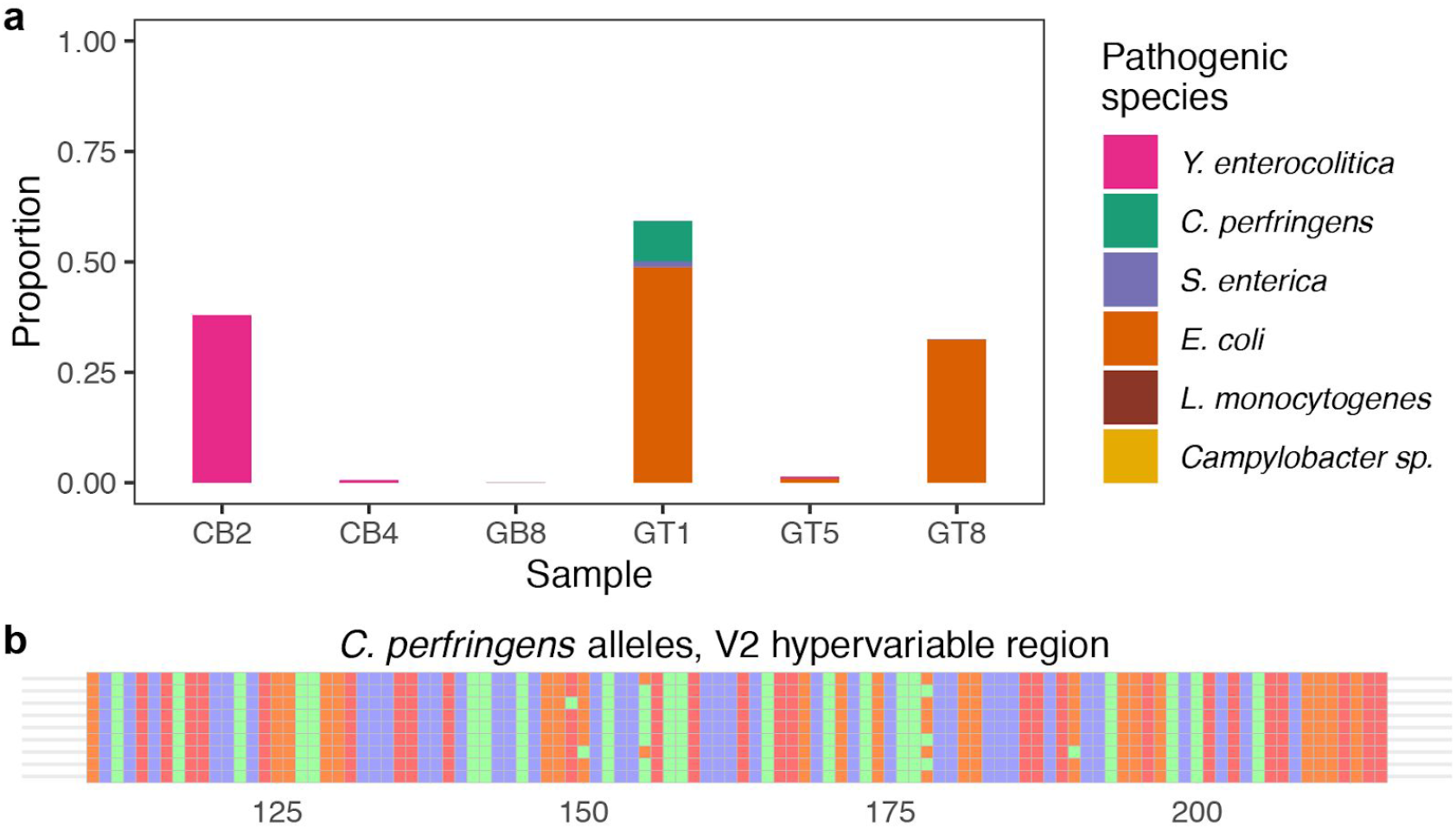
Foodborne pathogens detected in retail meat rinse samples. a) The species and relative abundance of six common foodborne pathogen species were determined from full-length LoopSeq 16S sequencing of six retail meat samples. Campylobacter and Listeria monocytogenes do not appear in these samples. b) Nine distinct alleles of the 16S rRNA gene were detected in the C. perfringens strain present in sample GT1. The first allele was present at roughly twice the abundance of the others, consistent with it having two copies in the genome while the rest have only one copy. C. perfringens has 10 copies of the 16S rRNA gene in its genome. Only the V2 region is shown for visual simplicity.

Most bacteria have multiple ribosomal operons, and the complete set of 16S rRNA gene allelic variation of many of the abundant foodborne pathogen strains present in these retail meat samples was fully resolvable (e.g. Figure 6b). However, in samples where multiple related strains of the same species were present, as was the case for *E. coli* in samples GT 1 and GT8, it was not always possible to unambiguously separate ASVs into strain-level bins, and the full allelic complement could not be captured for low abundance strains. Using a previously described strategy (Callahan 2019), we used the full complement of 16S alleles to determine that the *S. enterica* strain in sample GT 1 most closely matched a previously sequenced genome from the pathogenic Newport serovar, warranting further investigation into its potential pathogenicity. This direct inference could be made possible in the future with a better understanding of the phylogenetic coherence of the Newport serovar, and a more complete catalog of high-quality *Salmonella* genomes from pathogenic and non-pathogenic strains.

## Discussion

Amplicon sequencing is a cornerstone method in the life sciences, notably used for the characterization of microbial diversity in complex samples. The combination of PCR amplification and subsequent sequencing massively enriches a targeted genetic locus and provides detailed information about the genetic diversity at that locus. The fundamental constraint on amplicon sequencing until recently has been the short read lengths of modern high-throughput sequencers, but that constraint has been overcome by the rise of long-read sequencing technologies. In a world of expanding sequencing options, it is critical to understand the accuracy and economics of long-read amplicon sequencing technologies in order to know when each specific tool is the right choice for a given application.

Commercially available Synthetic Long-Read (SLR) sequencing technologies --- in which long sequencing reads are reconstructed from short reads containing molecular tags indicating origination from a common DNA molecule --- have been around for a number of years, but not for amplicon sequencing where homology between sequences is high. The LoopSeq technology recently commercialized by Loop Genomics applies unique molecular identifiers (UMIs) to each DNA molecule, and allows precise and accurate long-reads to be constructed from amplicon sequencing data. Here, we showed that LoopSeq amplicon sequencing attains higher accuracy than previously reported for commonly-used amplicon sequencing technologies, can scale out to sequence lengths of at least 6 kilobases while maintaining very high accuracy, and can be used to precisely survey the composition of complex microbial communities.

Accuracy is essential to many amplicon sequencing applications, and the level of accuracy achieved by LoopSeq may open up new opportunities. Long-read amplicon sequencing using the PacBio and Oxford Nanopore technologies (Callahan 2019; Eren 2019; Karst 2020) has received increased recent attention, with encouraging results demonstrating that per-base accuracy exceeding common short-read approaches can be obtained by combining long-read sequencing with molecular methods such as the construction of PacBio circular consensus (CCS) reads and appropriate bioinformatics. In the bacterial profiling application, multiple studies have shown that substantial improvements in species and subspecies resolution can be achieved by sequencing the entire ~1.5 kilobase 16S gene, rather than just segments of 100-500 bases as is most commonly practiced today, and that even greater resolution is achievable by extending the sequenced region to most or all of the *rrn* operon (Martijn 2019).

The new frontier of amplicon sequencing beyond 1.5 kilobases may be where the exceptional accuracy attainable by LoopSeq is most important. Using the manufacturer-recommended coverage thresholds and a common bioinformatics workflow, we showed here that >90% of LoopSeq full-length 16S reads were error free as compared to ~50% error-free full-length 16S PacBio CCS reads (Callahan 2019). 50% is a sufficiently high fraction of error-free reads for modern denoising methods to achieve single-nucleotide resolution with high accuracy, and thus the total community compositions determined after denoising full-length 16S rRNA gene sequences obtained using LoopSeq reads and PacBio CCS reads from the same human fecal samples were highly similar (Figure 4). However, if we consider amplicon sequencing of the entire ~5Kb *rrn* operon, then LoopSeq produces >50% error-free reads, while only ~10% of PacBio CCS reads would be expected to be error-free, significantly degrading the detection and resolution of lower-abundance members of a sampled community. Single-nucleotide resolution with high accuracy from the entire *rrn* operon opens the door for population-level analysis of genetic diversity, rather than just comparisons amongst species, and would significantly enhance discrimination and identification of critical sub-species variation such as pathogen and non-pathogen clades, especially in sample types where alternative shotgun metagenomics methods are challenged by large amounts of non-target DNA.

PCR amplicon chimeras, in which sequencing reads are produced that are a combination of multiple true DNA molecules, can be particularly pernicious for the accurate reconstruction of complex microbial communities. LoopSeq SLRs have a significant advantage over standard long-read amplicon sequencing approaches in this regard because LoopSeq SLRs are assembled from a consensus of UMI-tagged reads. As a result, chimeric molecules do not contribute to the consensus assembly unless the chimera formed at the first cycle or two of PCR and thus chimeric reads constitute a majority of the short reads for that UMI. Fortuitously, chimera rates are also at their lowest during early PCR cycles, further reducing the effective LoopSeq chimera rate. In standard amplicon sequencing technologies, chimeric molecules formed during all PCR cycles will be present in the final data. We note that this potential to leverage UMIs to identify and remove SLR chimeras was also demonstrated previously in a different SLR technology (Burke 2016).

We identified and described an SLR-specific structural sequencing error, the introgression, in which a long SLR is formed with an internal insertion of a short segment from another DNA molecule (Figure 2A). Introgressions are caused by the stochastic preponderance of short reads from chimeric DNA molecules over short regions of the SLR. Fortunately, the rate of introgressions in LoopSeq data is low, and no introgressions were detected when typical denoising of the Zymo mock community was performed that screened out singletons. Standard quality filtering based on expected errors further reduced the fraction of introgressions in the raw data (Figure 2). As LoopSeq or other SLR sequencing technologies become more widely used it may be useful to revisit the quality filtering approaches used for such data. Dips in the quality scores corresponding to introgressed LoopSeq regions suggest that moving window quality screens may be a useful tool for such data.

In this manuscript we focused on comparing LoopSeq to the long-read sequencing technologies developed by PacBio and Oxford Nanopore (ONT), because those technologies are currently the most widely used and are already commercially available. However, an important current research direction is the marriage of UMI methods with those long-read sequencing technologies to improve their accuracy. In a recent preprint, Karst and colleagues describe and evaluate such a method, and report achieving accuracy comparable to that reported here by pairing UMIs with Oxford Nanopore sequencing, and even higher levels of accuracy from pairing UMIs and PacBio CCS sequencing (Karst 2020). We expect this general approach, if not the exact implementation in Karst *et al.*, will prove to be an attractive option for highly-accurate long-read amplicon sequencing as it continues to develop. Each of these high-fidelity long-read sequencing methods (LoopSeq SLRs, PacBio + UMIs, ONT + UMIs) are able to achieve very high accuracy (>99.99% per-base) that can be increased further at the expense of throughput by increasing the number of reads per UMI (with the exception that ONT may experience an accuracy plateau for ONT due to continuing issues with systematic error modes (Karst 2020)). It is likely that in the next few years different highly-accurate long-read technologies will find unique niches within microbial genomics. For example, ONT sequencing might be advantageous for field applications of microbial genomics in which rapid diagnosis is key while LoopSeq and PacBio long reads might be more useful for applications that require high accuracy and that can be obtained within days, not hours. Perhaps the most important differentiating factor in the long-term will be cost. The future evolution of basal error rates and costs of the underlying short-read, PacBio and ONT sequencing technologies is unclear, but the most impactful high-fidelity long-read sequencing method will probably be determined by the evolution of cost efficiency and long-term choices about the value of sequencing accuracy.

The rapidly increasing accuracy and length of available amplicon sequencing technologies has laid bare the limitations of commonly used taxonomic assignment methods for 16S rRNA gene data. There are fundamental limits on the taxonomic resolution available from any marker-gene sequencing approach, but the most widely used methods for taxonomic assignment from 16S sequences were developed for short-read data and often do not even attempt to make taxonomic assignments beyond the genus level. Long-read amplicon sequencing data of the accuracy achieved by LoopSeq allows for species-level assignment from 16S rRNA gene data in most cases. Even higher levels of sub-species resolution are achievable, but substantial roadblocks exist in practice due to the multi-copy nature of the *rrn* operon in bacteria (Klappenbach 2001). Full-length 16S rRNA gene sequencing with single-nucleotide resolution will resolve all intragenomic variation between the 16S alleles carried by a single bacterium (Callahan 2019; Johnson 2019), but there is no currently automated way to reconstruct the bins containing alleles arising from a common genome. There is also no universal database that contains and labels the full complement of 16S rRNA gene alleles arising from each strain. Furthermore, many reference genomes created from short-read sequencing data do not resolve the multiple copies of the *rrn* operon at all. Preliminary evidence suggests that, at least in some cases, pathogenic and non-pathogenic *E. coli* can be distinguished from full-length 16S sequencing alone (Callahan 2019), but current taxonomic assignment methods do not approach that level of resolution.

There is a much larger potential universe of applications for LoopSeq and other highly accurate long-read sequencing technologies beyond profiling microbial communities by sequencing the *rrn* operon. One topical example is viral genomic sequencing, such as that being applied to the population genetics and genomic epidemiology of the SARS-nCov2 virus. Viral DNA exists as a tiny minority of the DNA in clinical samples, and standard SARS-nCov2 sequencing approaches rely on amplifying nearly 100 different genetic regions that are then stitched together to reconstruct a consensus genome sequence (DNA 2020). Highly-accurate long-read amplicon sequencing opens the door to simpler protocols using far fewer primers, and that can also achieve long-range linkage information (Gonzalez-Reiche 2020; Böhmer 2020; Sorensen 2020). These same advantages in simplicity, accuracy and long-range linkage information are driving the adoption of highly-accurate long-read amplicon sequencing for the study of HIV (Caskey 2017; Pauthner 2019), HLA/MHC (Westbrook 2015; Karl 2017; Shortreed 2020), and oncogene diversity in solid tumours (Chen 2018).

## Conclusion

Three aspects of microbial genomics will have significant bearing on bringing about a future of precision microbiology: the (1) accuracy of reading microbial genomes, the (2) discriminatory power of microbial sequencing reads, largely determined by read lengths, and the (3) quality of microbial sequence databases. Improvements in accuracy and length will feed directly into building better databases and generate a positive feedback loop that will eventually trivialize microbial identification and characterization. In this manuscript we showed how short-read sequencers can be used to generate microbial DNA reads with a combination of length and accuracy that matches and surpasses currently available methods. This LoopSeq technology leverages already widely available short read sequencers and is commercially-supported, a combination of attributes that could accelerate the uptake of accurate long-read sequencing in general. Amplicon sequencing that is an order-of-magnitude longer and an order-of-magnitude more accurate than the Illumina short-read standard is available today. We look forward to the ways this technology will be applied in the future.

## Methods

### Samples and DNA Extraction

Zymo mock community: The ZymoBIOMICS™ Microbial Community DNA Standard (P/N: D6306, Lot ZRC190811) was obtained from the manufacturer Zymo Research (Irvine, CA). The Zymo mock community contains genomic DNA from eight phylogenetically diverse bacteria, and two yeast strains not amplified by our 16S rRNA gene amplicon sequencing protocol. Note that five strains in ZymoBIOMICS™ standards were replaced with similar strains in Lot ZRC190633. The sample analyzed here is from a post-replacement lot.

Fungal isolates: Extracted and purified genomic DNA from the following six fungal isolates were obtained from the ATCC: *Saccharomyces cerevisiae Meyen ex E.C. Hansen* (Catalog Number ATCC 201389D-5), *Aspergillus oryzae var. oryzae* (Catalog Number ATCC 42149D-2), *Candida albicans (Robin) Berkhout* (Catalog Number ATCC10231D-5), *Trichoderma reesei Simmons* (Catalog Number ATCC 13631D-2), *Kluyveromyces lactis (Dombrowski) van der Walt (Catalog* Number ATCC 8585D-5) and *Penicillium chrysogenum Thom* (Catalog Number ATCC 10106D-2).

Bacterial isolates: Extracted and purified genomic DNA for the following three bacterial isolates were obtained from the ATCC: *Nitrosomonas europaea* (Catalog Number *ATCC 19718D-5):* https://www.ncbi.nlm.nih.gov/nuccore/AL954747, *Desulfovibrio desulfuricans* (Catalog Number *ATCC 27774D-5)*: https://www.ncbi.nlm.nih.gov/nuccore/CP001358, and *Salinispora tropica* (Catalog Number ATCC *CNB-440D-5):* https://www.ncbi.nlm.nih.gov/nuccore/CP000667.

Human fecal samples: Genomic DNA were obtained from three human fecal samples previously analyzed in a publication on PacBio long-read amplicon sequencing (Callahan 2019). The aliquots of DNA analyzed here were extracted as part of, and as described in, that publication.

Retail meat rinse samples: 250 mls buffered peptone water (BPW) were added to > 50 grams of retail meat (bone-in, skin on chicken breast; ground turkey, ground beef <85% lean; or bone-in pork chop), and samples were shaken at room temperature for 15 minutes at 250 rpm. Meat samples were incubated 18 hours in BPW at 37° C. 2ml rinsate was centrifuged at 1000g for 5 minutes, and supernatant was discarded. DNA was isolated from pellets using Lucigen MasterPure™ Gram Positive DNA Purification Kit according to manufacturer protocols. DNA was quantified by Qubit dsDNA HS Assay Kit assessed for purity using a Nanodrop 2000c (ThermoFisher Scientific).

### Sequencing library preparation

Sequencing of all sample was carried out from extracted genomic DNA that was made into sequence-ready libraries with commercially available LoopSeq long-read kits from Loop Genomics (protocols available at loopgenomics.com), and an informatics pipeline that converts short-reads into synthetic long reads (SLRs). The process involves attaching two DNA tags: one Unique Molecular Identifier (UMI) to each unique “parent” molecule and one sample-specific tag (i.e. a Sample Index) equally to all molecules in the same sample. Barcoded molecules are amplified, multiplexed and each UMI is distributed intramolecularly to a random position within each parent molecule. Molecules are then fragmented into smaller units at the position of each UMI, creating a library of UMI-tagged fragments with an average length of 400bp compatible with an Illumina sequencing platform run in PE150 mode.

Full-length 16S sequencing: For each LoopSeq Microbiome 16S kit, up to 24 samples were processed in multiplex and ~12,000 1.5kb molecules were sequenced per sample (~300k molecules from a complete kit run). 100-150M PE150 reads (50-75M clusters passing filter) were used for each sequencing run, yielding ~20 gigabases (Gb) of data. The complete sample preparation and sequencing protocol can be found in <u>this link</u>.

Full-length 18S-ITS sequencing: For each LoopSeq Mycobiome 18S-ITS kit, up to 24 samples were processed in multiplex and ~12,500 ~2.3kb molecules were sequenced per sample (~300k molecules from a complete kit run). 175-250M PE150 reads (87.5-125M clusters passing filter) were used for each sequencing run, yielding ~35 gigabases (Gb) of data. The complete sample preparation and sequencing protocol with sequencing instructions can be found in <u>this link</u>.

Bacterial whole genome sequencing: For each LoopSeq Bacterial Genome kit, up to 8 samples were processed in multiplex and ~40,000 ~5kb molecules were sequenced per sample (~320k molecules per library). 320M PE150 reads (160M clusters passing filter) were used for each sequencing run, yielding ~50 gigabases (Gb) of data. The complete sample preparation and sequencing protocol with sequencing instructions can be found in <u>this link</u>.

### Short read coverage of LoopSeq synthetic long reads (SLRs)

In general, greater short-read coverage of each SLR will result in a higher fraction of complete SLRs (i.e. SLRs that span the full targeted amplicon) and a lower error rate. Here we evaluated LoopSeq data with an average of 300 150bp reads per full-length 16S read (30x coverage), which is the recommended short-read coverage in the manufacturer protocols. The minimal number of 150bp short reads required to assemble a full-length 1.5kb 16S rRNA gene using the LoopSeq SPADES workflow is 30 (3x coverage), but this would result in a significantly higher per-base error rate. Correspondingly, higher than 30x short-read coverage per SLR would be expected to produce per-base error rates even lower than those reported here.

### Assembly of SLRs

Loop Genomics maintains a cloud-based platform for processing raw short-reads prepared with a LoopSeq kit into assembled SLR contigs. Within this pipeline, short-reads are trimmed using Trimmomatic (Bolger 2014) to remove adapter sequences before they are de-multiplexed based on their Loop Sample Index, which groups them by the sample from which they originated. Within a grouped sample, short-reads are next binned by UMI such that those with the same UMI are processed collectively through SPADES (Bankevich 2012). Reads sharing the same UMI are derived from the same original molecule, with each read covering a different region of the sequence. With enough short-read data to cover the full length of a long DNA molecule, it is possible to assemble the original long DNA molecule by linking overlapping short-reads through their shared sequence, and then arranging the reads in the correct order to rebuild the original 16S/18S-ITS/Genomic molecule sequence. Assembly attempts with fewer reads result in shorter SLRs with lower accuracy.

### Amplicon bioinformatics

The raw LoopSeq synthetic long reads (SLRs) were subjected to further quality filtering, denoising and chimera removal using the dada2 R package, largely following the long-read workflow previously established for PacBio long-read amplicon sequencing (Callahan 2019). Briefly, SLRs were screened for the presence of both forward and reverse primers of the full length 16S gene, and truncated to the region between those primers. Primer-free sequences were filtered based on the total expected errors (Flvyberg and Edgar, 2015). The relationship between the quality scores and the error rates was learned from the data, and the denoised amplicon sequence variants (ASVs) were inferred using the DADA2 algorithm (Callahan 2016).

Standard data processing used the default parameters for long-reads described in (Callahan 2019). High-sensitivity parameters appropriate for LoopSeq data were also developed and used in several analyses. The key differences between the high-sensitivity and default parameters were that the probability threshold for detecting new ASVs was made less stringent (OMEGA_A=1e-10), and the option to directly detect singleton sequences was enabled (DETECT_SINGLETONS=TRUE).

### Characterizing synthetic long-reads by error type

After screening for and removing primers, a moving window comparison was made between every synthetic long-read (SLR) present in the Zymo mock community data and the 27 “reference” sequences corresponding to the unique full-length 16S rRNA gene alleles present in each mock community strain. For each window of 50 nts (step size of 10 nts) the most similar reference sequence(s) were recorded, and a strain-level assignment for that SLR-window was made if all of the most similar reference sequences belonged to the same strain. Otherwise no strain-level assignment was made for that window.

Two types of structural errors --- Chimeras and Introgressions --- were assigned based on the results of the moving window comparison. A chimera assignment was made if the best match to the left-hand side and the right-hand side of the SLR were from different strains. An introgression assignment was made if an internal segment was assigned to a different strain than the rest of the SLR. Windows in which no strain assignment was made were ignored. More complex patterns (e.g. SLRs that contained windows assigned to three or more different strains) were designated as Complex errors. Finally, SLRs that had a consistent strain-level assignment throughout, but that had up to three mismatches with the closest reference sequence, were assigned as point errors.

### Assessing conserved/variable status of substitutions

The DADA2 algorithm provides a complete description of the nucleotides differences that distinguish each ASV from the “sibling” ASV from which it was divided. The ssu-align method (http://eddylab.org/software/ssu-align/) was used to align each “sibling” ASV to a model for the bacterial 16S rRNA gene, and thereby to classify each nucleotide in the “sibling” ASV as either conserved or variable. Together, these two sources of information allowed the distinguishing substitutions for each ASV identified by DADA2 to be classified as occurring in conserved or variable positions. The null expectation of a 4:1 conserved:variable ratio in the case of random sequencing error was determined by randomizing the positions of distinguishing substitutions.

### Species assignment for foodborne pathogens

Species-level assignments were made for all ASVs in the meat samples with genus-level assignments corresponding to the six foodborne pathogens of interest by taking a consensus of BLAST results. In short, relevant sequences were BLAST-ed against nt (excluding uncultured/environmental accessions). The species designations of all BLAST hits sharing the top score were collated. If all top-hit species designations agreed, a species assignment was made. Accessions with no species designation were ignored.

### Code Availability and Reproducible Analysis

Rmarkdown code to reproduce the results described in this paper is available at https://github.com/beniineb/LoopManuscript

## Data Availability

All sequencing data are available from the SRA under BioProject Accession PRJNA644197. Rmarkdown code to reproduce the results described in this paper is available at https://github.com/benjjneb/LoopManuscript

## Supplementary Figures

**Figure S1:**
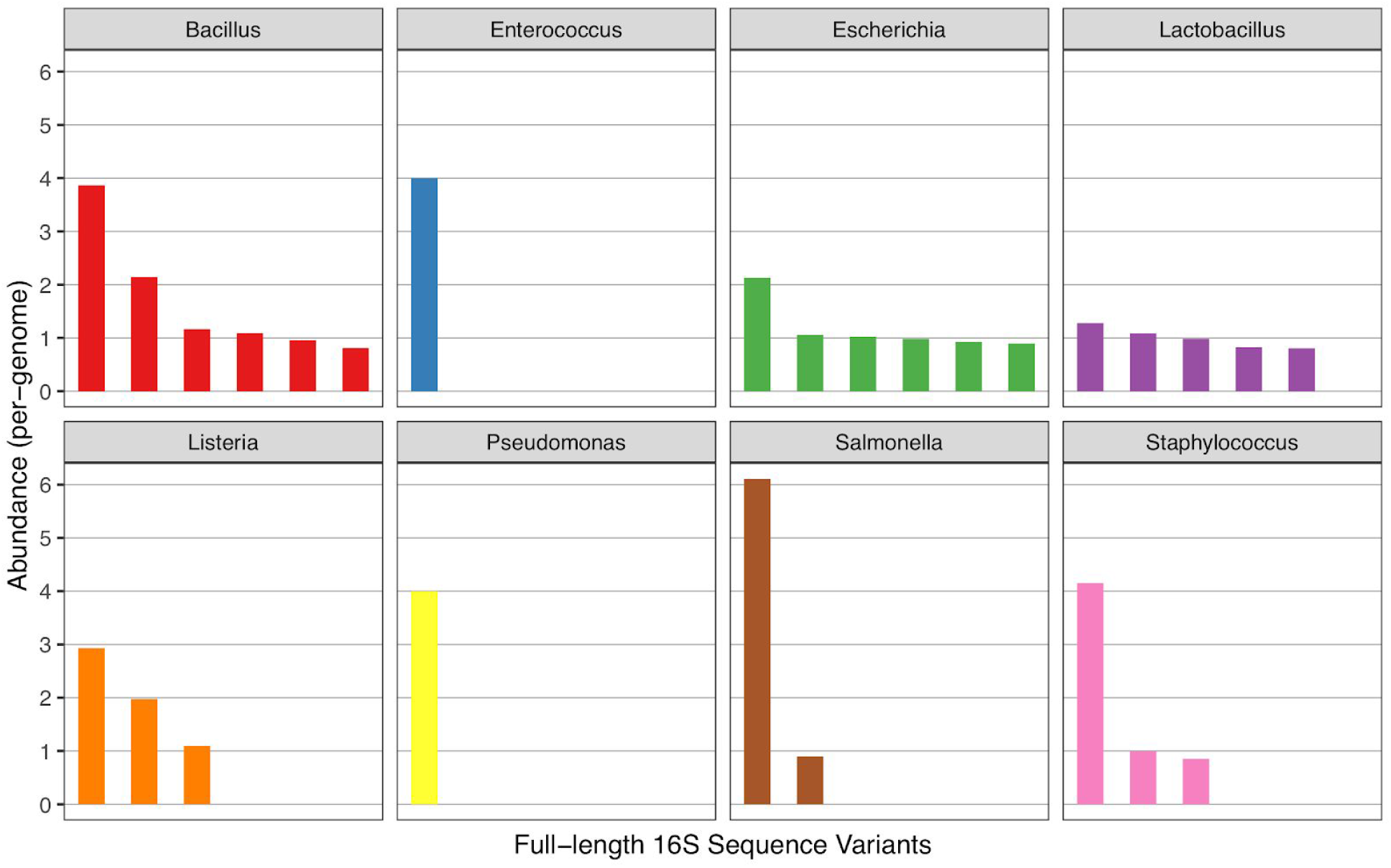
The abundances of all ASVs identified by LoopSeq and default DADA2 in the Zymo mock community. All abundances are scaled to the abundance of the corresponding genome in the amplified 16S rRNA gene data. The near-integer values of all genome-scaled abundances are consistent with each ASV representing a unique allele present in the multiple copies of the 16S rRNA gene present in the genomes of these strains.

